# Identification of viruses belonging to the family *Partitiviridae* from plant transcriptomes

**DOI:** 10.1101/2020.03.11.988063

**Authors:** Yeonhwa Jo, Won Kyong Cho

**Affiliations:** Research Institute of Agriculture and Life Sciences, College of Agriculture and Life Sciences, Seoul National University, Seoul 08826, Republic of Korea

**Keywords:** Viruses, *Partitiviridae*, Identification, Plant transcriptome

## Abstract

Viruses in the family *Partitiviridae* consist of non-enveloped viruses with bisegmented double-stranded RNA genomes. Viruses in this family have been identified from plants and fungi. In this study, we identified several viruses belonging to the family *Partitiviridae* using plant transcriptomes. From 11 different plant species, we identified a total of 74 RNA segments representing 23 partitiviruses. Of 74 RNA segments, 28 RNA segments encode RNA-dependent RNA polymerases (RdRp) while 46 RNA segments encode coat proteins (CPs). According to ICTV demarcation for the family *Partitiviridae*, 25 RNAs encoding RdRp and 41 RNAs encoding CP were novel RNA segments. In addition, we identified eight RNA segments (three for RdRp and five for CP) belonging to the known partitivruses. Taken together, this study provides the largest number of partitiviruses from plant transcriptomes in a single study.

## Introduction

Viruses in the family *Partitiviridae* consist of non-enveloped viruses with bisegmented double-stranded (ds) RNA genomes [1]. Each RNA segment of the viruses in the family *Partitiviridae* encodes a single protein including RNA-dependent RNA polymerase (RdRp) or coat protein (CP). Members in the family *Partitiviridae* can be divided into five different genera according to the host. For example, viruses in the genera *Alphapartitivirus* and *Betapartitivirus* are identified from either plants or fungi while viruses in the genus *Gammapartitivirus* and *Deltapartitivirus* are derived from only fungi and plants, respectively [1]. In addition, viruses in the genus *Cryspovirus* are identified from protozoa. Transmission of partitiviruses is occurred by seeds (plants), cell division and sporogenesis (fungi), and oocytes (protozoa) [1]. To date, there are more than 45 partitivirus species. In particular, there are five deltapartitivirus species infecting plants including *Pepper cryptic virus 1* (PCV-1) [2], *Pepper cryptic virus 2* (PCV-2), *Fig cryptic virus* (FCV) [3], *Beet cryptic virus 2* (BCV2), and *Beet cryptic virus 3* (BCV3) [4] according to International Committee on Taxonomy of Viruses (ICTV).

To identify novel partitiviruses with dsRNA genomes, dsRNA extraction followed by next-generation sequencing (NGS) is an efficient approach. Based on those approaches, several partitiviruses have been identified. For example, Melon necrotic spot virus was identified from watermelon plant (*Citrullus lanatus* Thunb) using SOLiD NGS analysis [5]. Pittosporum cryptic virus 1 was identified from an Italian pittosporum plant by NGS with extracted dsRNA [6]. Similarly, application of NGS to extracted dsRNA has resulted in identification of Trichoderma atroviride partitivirus 1 from the fungal species *Trichoderma atroviride* [7].

Moreover, instead of traditional molecular cloning methods, *in silico* data analyses has resulted in identification of diverse dsRNA viruses from expressed sequence tag (EST) database [8] and plant transcriptomes [9-13]. Moreover, 120 mycoviruses infecting fungi have been identified from fungal transcriptomes including several partitiviruses [14].

Recently, we carried out *in silico* data analyses to identify virus-associated contigs from plant transcriptomes. As a result, we identified a total of 74 RNA segments representing 23 partitiviruses belonging to the family *Partitiviridae* from 11 different plant species,

## Materials and methods

### *De novo* transcriptome assembly and BLAST search

We downloaded RNA-Sequencing (RNA-Seq) from the sequence read archive (SRA) database. The downloaded raw sequence data were *de novo* assembled by Trinity assembler with default parameters [15]. The assembled contigs (transcriptome) were blasted against the database containing all viral genomes for the family *Potyviridae*, which were manually collected, using TBLASTX with a cut-off E-value of 1e−3. After that, we extracted virus-associated contigs based on the TBLASTX result. To eliminate sequences derived from the plants and other contaminants, the identified virus-associated contigs were subjected to BLASTN search against NCBI’s nucleotide (NT) database. Finally, we obtained clean virus-associated contigs.

### Prediction of open reading frames

Of identified virus-associated contigs, the contigs more than 1 kb in length were selected for the search of open reading frames (ORFs) using ORFfinder program (https://www.ncbi.nlm.nih.gov/orffinder/). The identified ORF protein sequences were subjected to BLASTP search against non-redundant (NR) protein database with a cut-off E-value of 1e−3.

### Virus classification

Based on BLASTP results with individual viral ORF amino acid sequence, we determined virus taxonomy. The newly identified virus was named based on the identified virus host and the homologous viral genus. In case of partitiviruses, the RNA segment encoding RdRp was named as RNA1 while other RNA segment encoding CP was named RNA2. Based on ICTV’s demarcation for the family *Partitiviridae*, we determined novel viruses. For instance, amino acid sequence identity for RdRp and CP was less than 90% and 80%, respectively for species demarcation in the family *Partitiviridae*. For pairwise sequence alignment, we used sequence demarcation tool (SDT) program version 1.2 [16].

### Phylogenetic analyses

To reveal phylogenetic relationship of identified viruses, we constructed phylogenetic trees using MEGA7 program [17]. We retrieved top ten viral protein sequences homologous to identified viral proteins such as RdRp and CP. The obtained amino acid sequences with the identified viral protein were together aligned by ClustalW program with default parameter implemented in MEGA7 program. The aligned protein sequences were subjected to construction of phylogenetic tree. The phylogenetic tree was inferred by using the Maximum Likelihood method based on the JTT matrix-based model with with a bootstrap of 100 replicates [18].

## Results

### Identification of Alloteropsis cryptic virus 1 and 2 from *Alloteropsis semialata*

We identified six contigs associated with partitiviruses from the transcriptome of *Alloteropsis semialata* (BioProject PRJNA310121), which is a perennial grasses in the family *Poaceae* [19].

The sizes of identified contigs ranged from 1,326 bp to 1,712 bp. The six contigs encode a single ORF. Of them, two contigs encode RdRp while four contigs encode CP. Based on BLASTP search and the identified host, two novel viruses were tentatively named Alloteropsis cryptic virus 1 (AlCV1) and Alloteropsis cryptic virus 2 (AlCV2). Both AlCV1 and AlCV2 consists of three RNA fragments: RNA1 (RdRp), RNA2 (CP), and RNA3 (CP). The RNA1 of AlCV1 showed sequence similarity to RdRp of Arhar cryptic virus-I (YP_009026407.1), which is an unclassified partitivirus, The RNA2 and RNA3 of AlCV1 showed sequence similarity to CPs of White clover cryptic virus 1 (YP_086755.1) [20] and Citrullus lanatus cryptic virus (APT68924.1) [21], respectively. The RNA1 of AlCV2 showed sequence similarity to RdRp of Raphanus sativus cryptic virus 3 (YP_002364401.1) [22] whereas RNA2 and RNA3 of AlCV2 showed sequence similarity to CPs of Arhar cryptic virus-I (YP_009026398.1 and YP_009026398.2), respectively. The phylogenetic tree based on RdRp amino acid sequences showed that AlCV1 RNA1 and AlCV2 RNA2 were different from each other. In addition, the RNA2 and RNA3 of AlCV1 in the same clade were different from those of AlCV2 in the same clade according to the phylogenetic tree using CP amino acid sequences.

### Identification of Amaranthus cryptic virus 1–4 from *Alloteropsis semialata*

We identified ten contigs associated with partitiviruses from the transcriptome of *Amaranthus tuberculatus* (BioProject PRJNA432348), which is a weed species in the family *Amaranthaceae* known as waterhemp [23]. The sizes of identified contigs ranged from 667 bp to 2,369 bp. The 10 contigs encode a single open reading frame (ORF). Of them, four contigs encode RNA dependent RNA polymerase (RdRp) while six contigs encode coat protein (CP). Based on BLASTP search and the identified host, the identified four novel viruses were tentatively named Amaranthus cryptic virus 1 (AmCV1), Amaranthus cryptic virus 2 (AmCV2), Amaranthus cryptic virus 3 (AmCV3), and Amaranthus cryptic virus 4 (AmCV4). RNA1 of AmCV1 showed sequence similarity to RdRp of Persimmon cryptic virus (YP_006390091.1) whereas RNA1 of Pepper cryptic virus 2 showed sequence similarity to RdRp of Pepper cryptic virus 2 (AVL84364.1). RNA2 of AmCV1 displayed sequence similarity to CP of Pepper cryptic virus 1 (AYA43792.1) whereas RNA2 of AmCV2 displayed sequence similarity to CP of Persimmon cryptic virus (YP_006390090.1). For AmCV3, we identified two sequences (isolates Won and Cho) of RNA1 and RNA2. Both AmCV3 RNA1 isolates showed sequence similarity to RdRp of Hop trefoil cryptic virus 2 (YP_007889825.1) whereas both AmCV3 RNA2 showed sequence similarity to CP of Red clover cryptic virus 1 (AWK57379.1). RNA1 of AmCV4 showed sequence similarity to RdRp of Hop trefoil cryptic virus 2 whereas RNA2 of AmCV4 showed sequence similarity to CP of Primula malacoides virus China/Mar2007 (YP_003104769.1) [24]. The phylogenetic tree using RdRp amino sequences for AmCV1 to AmCV4 demonstrated that two distinct groups. The first group contains RNA1 of AmCV1 and AmCV2 whereas the second group includes RNA1 of AmCV3 and AmCV4. Based on phylogenetic analysis, it is likely that RNA1 of AmCV4 is a partial sequence of RNA1 for AmCV3. In case of RNA2 of AmCV1 to AmCV4, all CP sequences are different from each other.

### Identification of Ambrosia cryptic virus 1–2 from *Ambrosia trifida*

We identified four contigs associated with partitiviruses from the transcriptome of *Ambrosia trifida*, which is a highly competitive annual weed known as giant ragweed (BioProject: PRJNA267208). The sizes of identified contigs ranged from 1,608 bp to 1,966 bp. The four contigs encode a single open reading frame (ORF). Of them, two contigs encode RdRp while other two contigs encode CP. Based on BLASTP search and the identified host, the newly identified two viruses were tentatively named Ambrosia cryptic virus 1 (AmbCV1) and Ambrosia cryptic virus 2 (AmbCV2), respectively. RNA1 of AmbCV1 showed sequence similarity to RdRp of Raphanus sativus cryptic virus 4 (ATG29853.1) [25] while RNA1 of AmbCV2 showed sequence similarity to RdRp of Pepper cryptic virus 2 (AVL84364.1). RNA2 of AmbCV1 and AmbCV2 showed sequence similarity to CPs of Red clover cryptic virus 1 (AVL84364.1) and White clover cryptic virus 1 (YP_086755.1), respectively. The phylogenetic tree for AmbCV1 and AmbCV2 using RdRp sequences showed that two viruses are clearly different from each other. In addition, the phylogenetic tree using CP sequences revealed that RNA2 of AmbCV1 were distinctly related with RNA2 of AmbCV2.

### Identification of Camellia cryptic virus 1 from *Camellia sinensis*

We identified three contigs associated with partitiviruses from the transcriptome of *Camellia sinensis*, which is an economically important tea plant (BioProject: PRJNA295355). Of them, BLASTP search with three contigs showed sequence similarity to camellia oleifera cryptic virus 1 (GenBank QBQ83754.1) [26], which is a member of the genus *Deltapartitivirus* in the family *Partitiviridae*. The newly identified virus was tentatively named Camellia cryptic virus 1 (CCV1) isolate Won. RNA1 (1,681 bp) of CCV1 encodes an ORF (477 aa) showing sequence similarity to RdRp of Camellia oleifera cryptic virus 1 (QBQ83753.1) with 100% coverage and 92.87% amino acid identity. RNA2 (1,318 bp) of CCV1 encodes an ORF (344 aa) showing sequence similarity to CP2 of camellia oleifera cryptic virus 1 (QBQ83755.1) with 100% coverage and 90.41% amino acid identity. RNA3 (1,410 bp) of CCV1 encodes an ORF (343 aa) showing sequence similarity to CP1 of Camellia oleifera cryptic virus 1 (QBQ83754.1) with 100% coverage and 90.96% amino acid identity. Based on BLASTP results, CCV1 is an isolate of Camellia oleifera cryptic virus 1 in the genus *Deltapartitivirus* in the family *Partitiviridae*.

### Identification of Camellia cryptic virus 2 from *Camellia reticulata*

We identified five contigs associated with partitiviruses from the transcriptome of *Camellia reticulate* (Bioproject: PRJNA297756), which is well known for its ornamental and high quality oil seeds belonging to the genus *Camellia* in the family *Theaceae* [27]. The newly identified virus was tentatively named Camellia cryptic virus 2 (CCV2) isolate Won. CCV2 consisted of five RNA fragments. RNA1 (1,510 bp) of CCV2 encodes an ORF (476 aa) showing sequence similarity to RdRp of Melon partitivirus (YP_009551627.1) with 99% coverage and 65.47% amino acid identity. RNA2 (1,719 bp) of CCV2 encodes an ORF (484 aa) showing sequence similarity to CP of Cucumis melo cryptic virus (QBC66122.1) [28] with 99% coverage and 36% amino acid identity. RNA3 (1,396 bp) of CCV2 encodes an ORF (343 aa) showing sequence similarity to CP1 of Camellia oleifera cryptic virus 1 (QBQ83754.1) [26] with 100% coverage and 91.25% amino acid identity. RNA4 (1,383 bp) of CCV2 encodes an ORF (344 aa) showing sequence similarity to CP2 of camellia oleifera cryptic virus 1 (QBQ83755.1) with 100% coverage and 89.24% amino acid identity. RNA5 (1,383 bp) of CCV2 encodes an ORF (396 aa) showing sequence similarity to CP of Clohesyomyces aquaticus partitivirus 1 (AZT88587.1) [29] with 100% coverage and 53.52% amino acid identity. The phylogenetic tree using RdRp amino acid sequences showed that CCV1 is closely related with the known Camellia oleifera cryptic virus 1; however, CCV2 in the same clade with Beet cryptic virus 3 was different from CCV1. The phylogenetic tree using CP amino acid sequences demonstrated that CCV1 RNA3 was closely related with COCV1 RNA2 whereas CCV1 RNA2 was closely related with COCV1 RNA3. However, RNA2 of CCV2 and RNA3 of CCV2 were different from each other. Therefore, CCV2 is a novel virus in the genus *Deltapartitivirus* in the family *Partitiviridae*.

### Identification of Dactylorhiza cryptic virus 1–3 from *Dactylorhiza incarnata*

We identified six virus-associated contigs associated with partitiviruses from the transcriptome of *Dactylorhiza incarnate* (BioProject PRJNA317244), which is a perennial species belonging to the family *Orchidaceae* [30]. The sizes of contigs ranged from 1,333 bp to 1,590 bp. Each contig encodes an ORF. Three contigs encode RdRp while other three contigs encode CP. The three identified viruses were tentatively named Dactylorhiza cryptic virus 1 (DCV1), Dactylorhiza cryptic virus 2 (DCV2), and Dactylorhiza cryptic virus 3 (DCV3). RNA1 of DCV1 showed sequence similarity to RdRp of Beet cryptic virus 2 (QCF59322.1). RNA2 of DCV2 showed sequence similarity to CP of Persimmon cryptic virus (YP_006390090.1). RNA1 and RNA2 of DCV2 showed sequence similarity to RdRp of Persimmon cryptic virus (YP_006390091.1) and CP of Pepper cryptic virus 1 (BAV93049.1), respectively. RNA1 and RNA2 of DCV3 showed sequence similarity to RdRp of Sinapis alba cryptic virus 1 (YP_009255398.1) and CP of Pittosporum cryptic virus-1 (CEJ95597.2), respectively. The two different phylogenetic trees using RdRp and CP amino acid sequences, respectively, revealed that DCV1–3 belong to the different groups of partitiviruses.

### Identification of Lomandra cryptic virus 1 from *Lomandra longifolia*

We identified three virus-associated contigs associated with partitiviruses from the transcriptome of *Lomandra longifolia*, which is an Australian native species resistant to *Phytophthora cinnamomi* (BioProject: PRJNA326999) [31]. Of them, BLASTP search with three contigs showed sequence similarity to Camellia oleifera cryptic virus 1 (GenBank QBQ83754.1) [26], which is a member of the genus *Deltapartitivirus* in the family *Partitiviridae*. The newly identified virus was tentatively named Lomandra cryptic virus 1 (LCV1) isolate Won. RNA1 (1,314 bp) of LCV1 encodes an ORF (363 aa) showing sequence similarity to RdRp of Camellia oleifera cryptic virus 1 (QBQ83753.1) with 100% coverage and 74.66% amino acid identity. RNA2 (1,350 bp) of LCV1 encodes an ORF (343 aa) showing sequence similarity to CP2 of Camellia oleifera cryptic virus 1 (QBQ83755.1) with 95% coverage and 38.60% amino acid identity. RNA3 (1,321 bp) of LCV1 encodes an ORF (345 aa) showing sequence similarity to CP1 of Camellia oleifera cryptic virus 1 (QBQ83754.1) with 98% coverage and 51.47% amino acid identity. Based on BLASTP result and phylogenetic analyses, LCV1 is a novel virus in the genus *Deltapartitivirus* in the family *Partitiviridae*. In addition, this is the first report of a partitivirus from *Lomandra longifolia*.

### Identification of Helianthus cryptic virus 1 from *Helianthus niveus*

We identified eight virus-associated contigs associated with partitiviruses from the transcriptome of *Helianthus niveus*, which is a diploid annual or perennial sunflower known as snowy sunflower in the family *Asteraceae* (BioProject: PRJNA320343) [1]. The sizes of identified contigs ranged from 1,384 bp to 2,399 bp. All contigs encode an ORF. Three contigs encode RdRp while other five contigs encode CP. The newly identified virus was named Helianthus cryptic virus 1 (HCV1). RNA1 of HCV1 shows sequence similarity to RdRp of Hop trefoil cryptic virus 2 (YP_007889825.1) while RNA2 of HCV1 shows sequence similarity to CP of Primula malacoides virus China/Mar2007 (YP_003104769.1). In addition, we identified two additional isolates: Cho and Kyong for RNA1 of HCV1 and four additional isolates: Cho, Kyong, Yeon, and Sik for RNA2 of HCV1. The phylogenetic trees using RdRp and CP amino acid sequences showed that all isolates for HCV1 were grouped together.

### Identification of Panax cryptic virus 1–4 *from Panax notoginseng*

We identified 15 contigs associated with partitiviruses from the two different transcriptomes of *Panax notoginseng*, which is a traditional Chinese medicine belonging to the genus *Panax* in the family *Araliaceae*. Nine contigs were identified from the first study (BioProject PRJNA393585) [32] and six contigs were identified from the second study (BioProject PRJNA472654) [33]. The sizes of identified contigs ranged from 1,013 bp to 1,836 bp. All contigs encode an ORF. From the first study, four different partitiviruses (Panax cryptic virus 1–4) (PCV1–4) were identified whereas three different partitiviruses (PCV1–3) were identified from the second study. The phylogenetic tree using RdRp amino acid sequences, RdRp and CP for PCV1–4 were different from each other. For example, RNA1 and RNA2 of PCV2 showed sequence similarity to RdRp (YP_006390091.1) and CP (YP_006390090.1) of Persimmon cryptic virus, respectively. Based on amino acid identity and phylogenetic analyses, we found that PCV1–3 in the second study were different isolates for PCV1–3 from the first study.

### Identification of Rhodiola cryptic virus 1–2 *from Rhodiola rosea*

We identified 11 contigs associated with partitiviruses from the transcriptome of *Rhodiola rosea*, which is a cold-tolerant perennial plant belonging to the family *Crassulaceae* used as traditional medicine (BioProject PRJNA398393) [34]. The sizes of 11 contigs ranged from 930 bp to 2,730 bp. Each contig encodes an ORF. Two contigs encode RdRp while other nine contigs encode CP. We identified two different partitiviruses were tentatively named Rhodiola cryptic virus 1 (RCV1) and Rhodiola cryptic virus 2 (RCV2). RCV1 consists of an RdRp and six CPs while RCV2 consists of an RdRp and five CPs. RNA1 of RCV1 showed sequence similarity to RdRp of Lysoka partiti-like virus (AWV67012.1) while RNA1 of RCV2 showed sequence similarity to RdRp of Medicago sativa deltapartitivirus 1 (ATJ00052.1). The phylogenetic tree using RdRp amino acid sequences showed that RCV1 and RCV2 belong to the different group of partitiviruses. The phylogenetic tree using CP amino acid sequences for RCV1 and RCV2 demonstrated that several groups of CPs. For example, RNA2 of RCV1 was closely related with Medicago sativa alphapartitivirus 1. Both RNA4 and RNA5 of RCV1 showed close genetic relationship with Lysoka partiti-like virus. RNA2–4 of RCV2 were clustered together with RNA6 of RCV1. In addition, RNA1 of RCV2 showed sequence similarity to CP of Raphanus sativus cryptic virus 1.

### Identification of Vicia cryptic virus *from Vicia faba*

We identified three contigs associated with partitiviruses from the transcriptome of *Vicia faba*, which is a commercially important nitrogen-fixing legume known as faba bean belonging to the family *Fabaceae* [35]. The sizes of three contigs ranged from 1,662 bp to 1,701 bp. Three contigs encode an ORF. Two contigs showed sequence similarity to RdRp (ABN71239.1) and CP (ABN71235.1) of known Vicia cryptic virus (VCV), respectively [36]. However, a contig (1,697 bp) showed sequence similarity to CP of Red clover cryptic virus 1. Based on the BLAST result, we named the identified virus as Vicia cryptic virus isolate Won. The phylogenetic tree using RdRp amino acid sequences demonstrated that RNA1 of VCV isolate Won was closely related with other isolates of VCV. However, the phylogenetic tree using CP amino acid sequences showed that RNA2 is a new isolate of VCV; however, RNA2 is newly identified RNA segment for VCV.

### Summary

In this study, we performed a large scale *in silico* data analyses to identify viruses belonging to the family *Partitiviridae*. We identified a total of 74 RNA segments associated with partitiviruses from 11 different plant species. Detailed information for identified RNA segment was provided in Table 1. We generated 20 different phylogenetic trees to reveal phylogenetic relationships of newly identified partitiviruses with known partitiviruses (Supplementary Figure). In addition, we provided BLASTP results of individual ORF for identified partitiviruses (Supplementary Table 1). Of the 11 plant species, Panax notoginseng (11 RNA segments representing four different partitiviruses) and Amaranthus tuberculatus (10 RNA segments representing four different partitiviruses) contain the largest number of RNA segments associated with partitiviruses (Table 2). Except eight RNA segments associated with Camellia oleifera cryptic virus 1 and Vicia cryptic virus, all RNA segments were novel. The phylogenetic tree using 25 RdRp amino acid sequences revealed four different groups of identified partitiviruses (Figure 1a) whereas the phylogenetic tree using 41 CP amino acid sequences showed five groups of partitiviruses. Taken together, this study provides the largest number of novel partitiviruses in a single study.

**Table 1.**
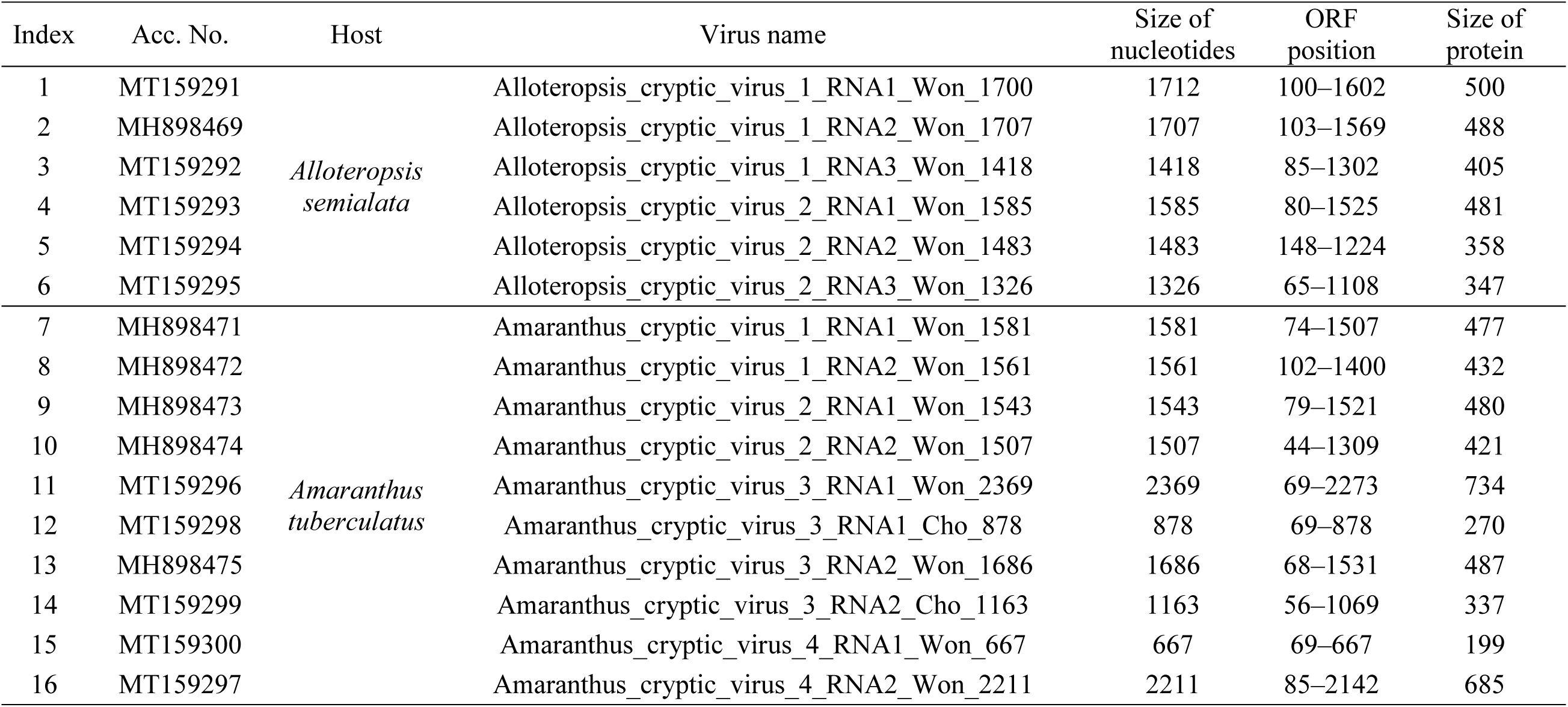
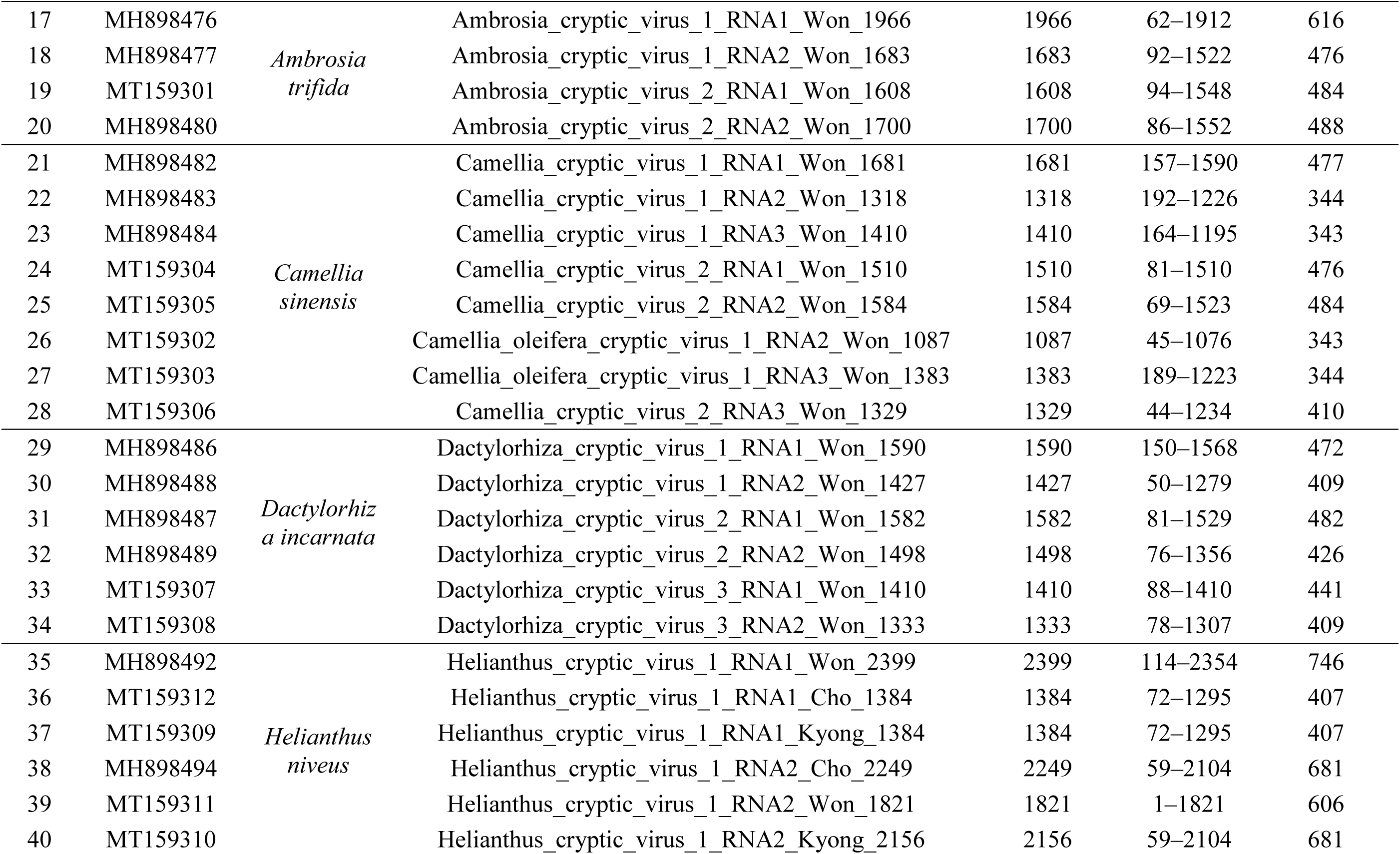
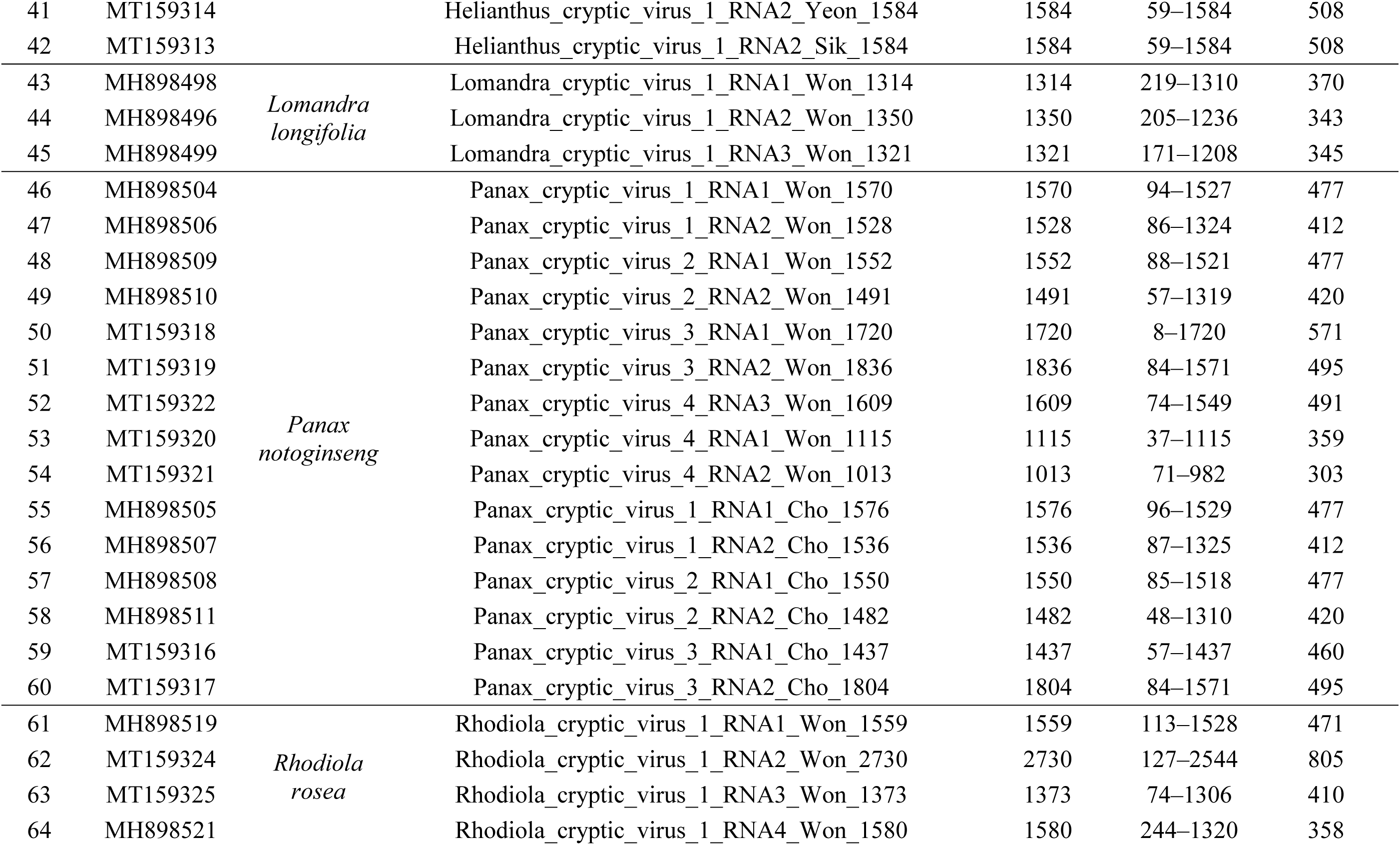
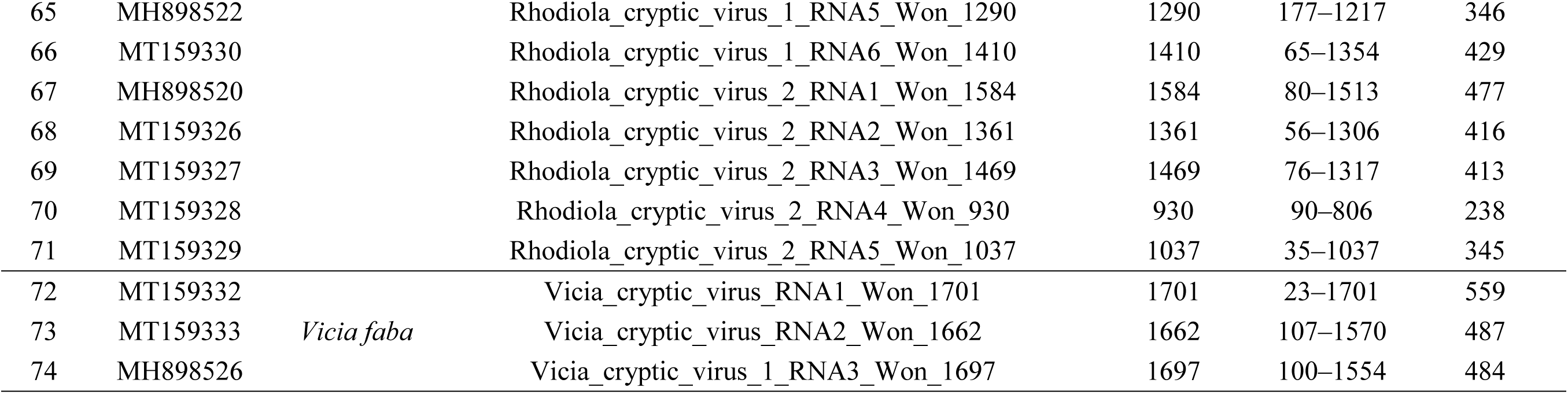
Detailed information for identified RNA segments associated with partitiviruses. Individual viral sequence was deposited in NCBI’s GenBank with respective accession number (Acc. No.). The name of plant host, virus name, size of nucleotide sequences, ORF position, and size of viral protein were provided.

**Table 2.** Summary of number of identified RNA segments, number of viruses, number of RdRp and number of CP in individual plant species.

| Plant species | No. of<br>RNA | No. of viruses | No. of<br>RdRp | No. of<br>CP |
| --- | --- | --- | --- | --- |
| *Alloteropsis semialata* | 6 | 2 | 2 | 4 |
| *Amaranthus tuberculatus* | 10 | 4 | 5 | 5 |
| *Ambrosia trifida* | 4 | 2 | 2 | 2 |
| *Camellia reticulata* | 5 | 2 | 1 | 4 |
| *Camellia sinensis* | 3 | 1 | 1 | 2 |
| *Dactylorhiza incarnata* | 6 | 3 | 3 | 3 |
| *Helianthus niveus* | 8 | 1 | 3 | 5 |
| *Lomandra longifolia* | 3 | 1 | 1 | 2 |
| *Panax notoginseng* | 15 | 4 | 7 | 8 |
| *Rhodiola rosea* | 11 | 2 | 2 | 9 |
| *Vicia faba* | 3 | 1 | 1 | 2 |

**Figure 1.**
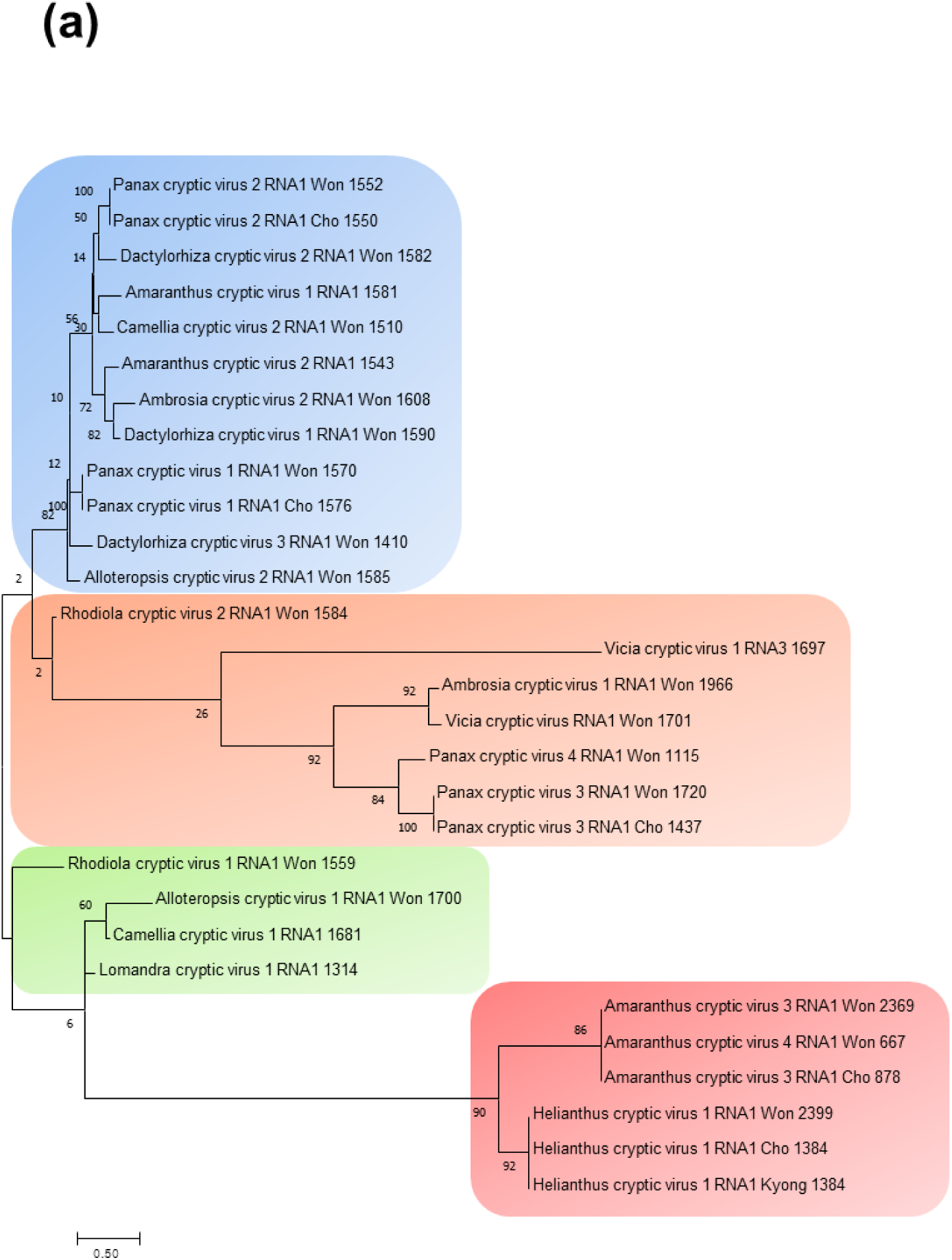
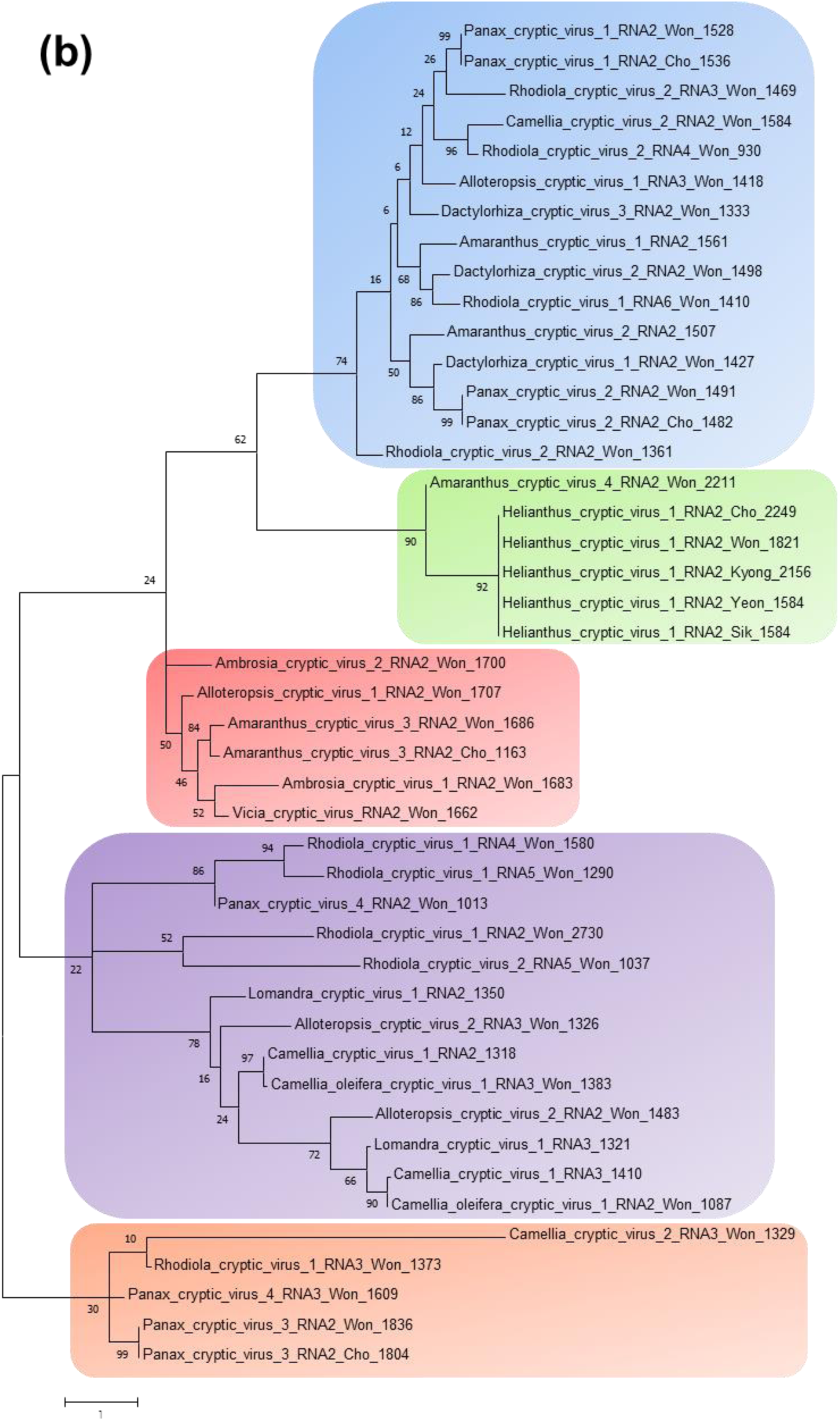
Phylogenetic relationships of identified viruses based on RdRp (a) and coat protein (CP) (b) amino acid sequences. The phylogenetic trees were constructed using the Maximum Likelihood method based on the JTT matrix-based model by MEGA 7.0 with a bootstrap of 100 replicates.

## Declarations

### Ethics approval and consent to participate

Not applicable

### Availability of data and materials

Viral genome sequences for identified viruses were deposited in GenBank with respective accession number provided in Table 1.

## Acknowledgements

This work was supported by a National Research Foundation of Korea (NRF) grant funded by the Korea government Ministry of Education (No. NRF-2018R1D1A1B07043597) and the support of the “Cooperative Research Program for Agriculture Science & Technology Development” (PJ01498301) conducted by the Rural Development Administration, Republic of Korea.

## Supplementary Figures

Phylogenetic relationships of identified viruses and other members of the family *Partitiviridae* based on RdRp and CP amino acid sequences. The phylogenetic trees were constructed using using the Maximum Likelihood method based on the JTT matrix-based model by MEGA 7.0 with a bootstrap of 100 replicates.The orange colored boxes indicate the identified partitiviruses from this study.

## Supplementary Table

**Table S1.** BLASTP results of identified viral proteins associated with partitiviruses.

The individual ORF proteins were subjected to BLASTP search against NCBI’s non redundant protein database. The best matched protein for individual viral protein with amino acid sequence coverage, identity, and accession number were provided.

