## Supplementary Figures for "Identification of viruses belonging to the family *Partitiviridae* from plant transcriptomes"

### Alloteropsis cryptic virus 1–2 using RdRp

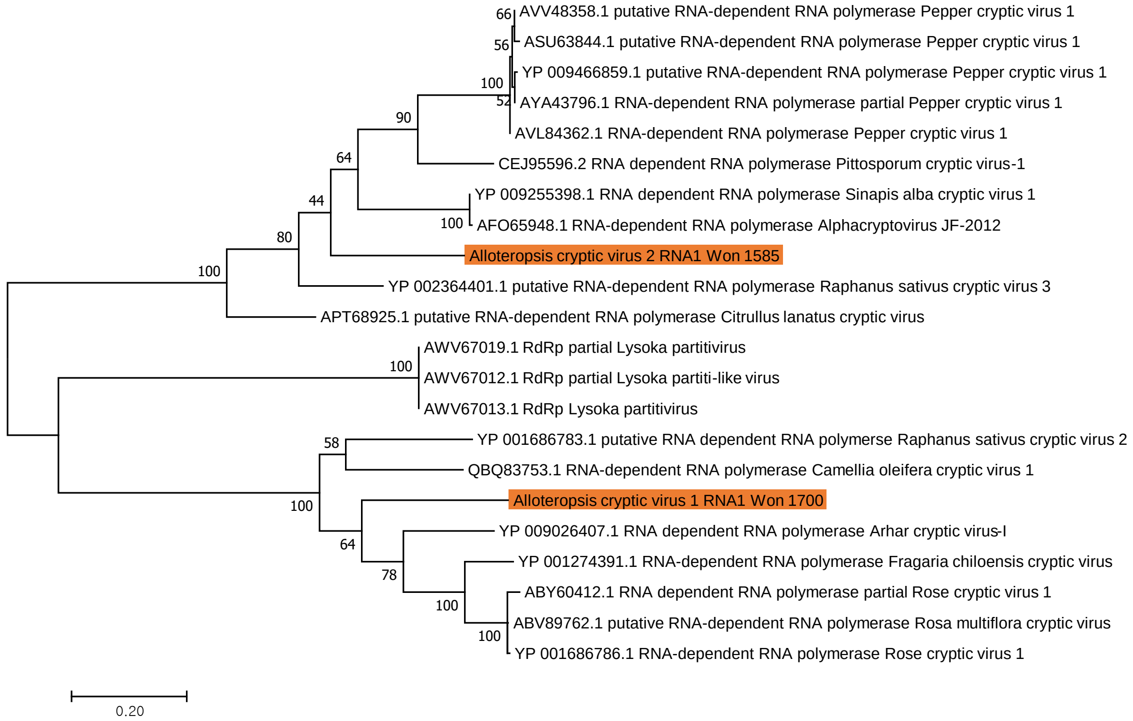

### Alloteropsis cryptic virus 1–2 using CP

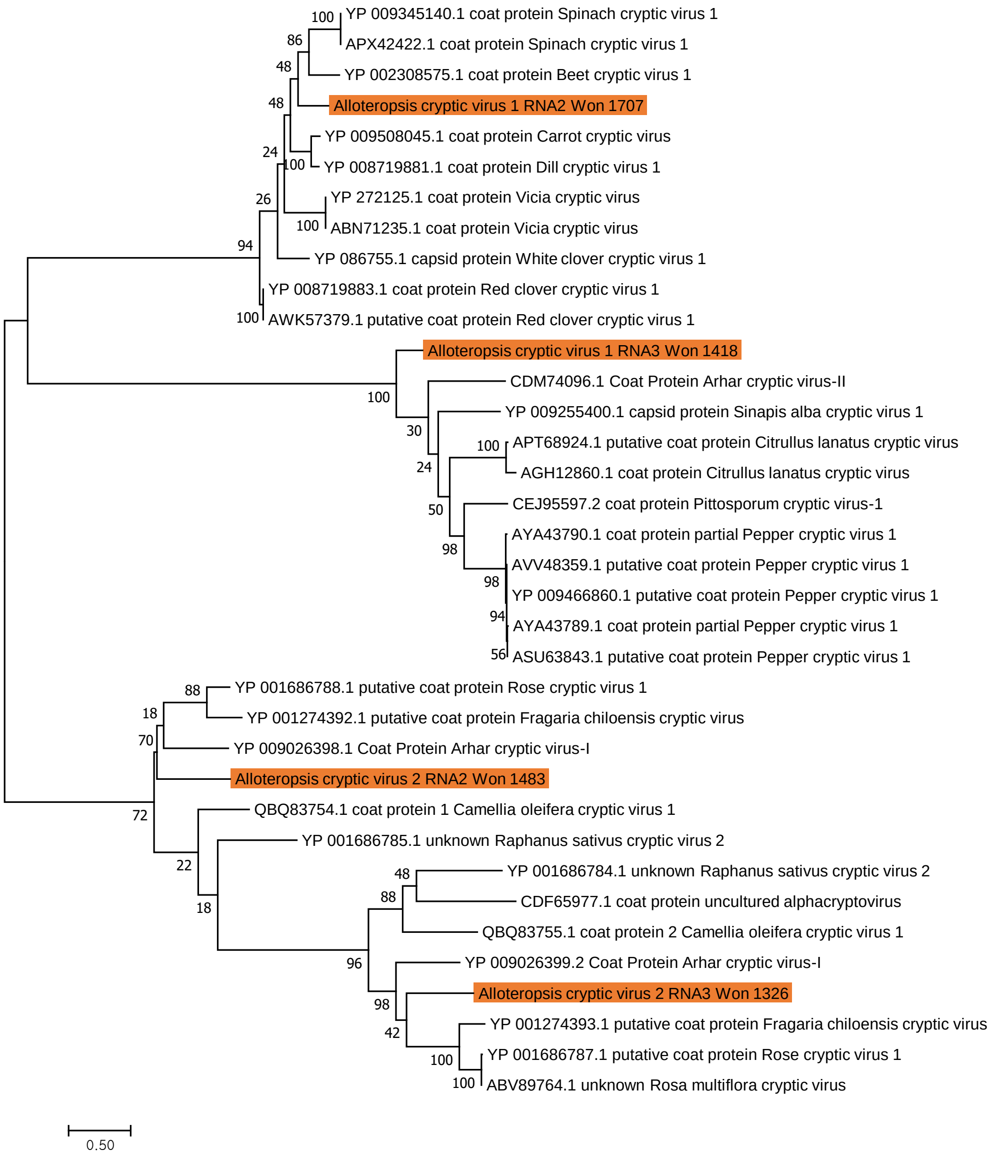

### Amaranthus cryptic virus 1–4 using RdRp

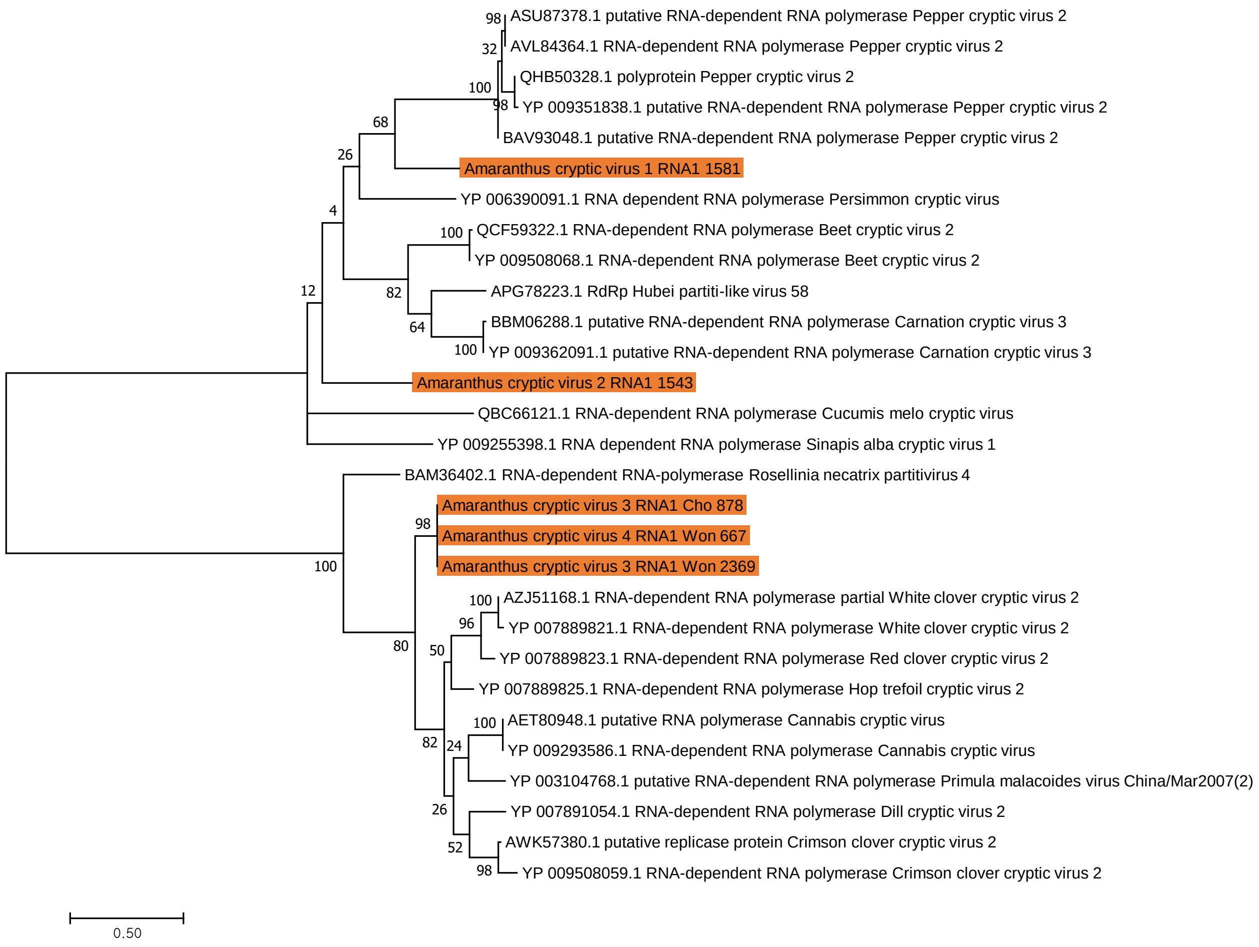

### Amaranthus cryptic virus 1–4 using CP

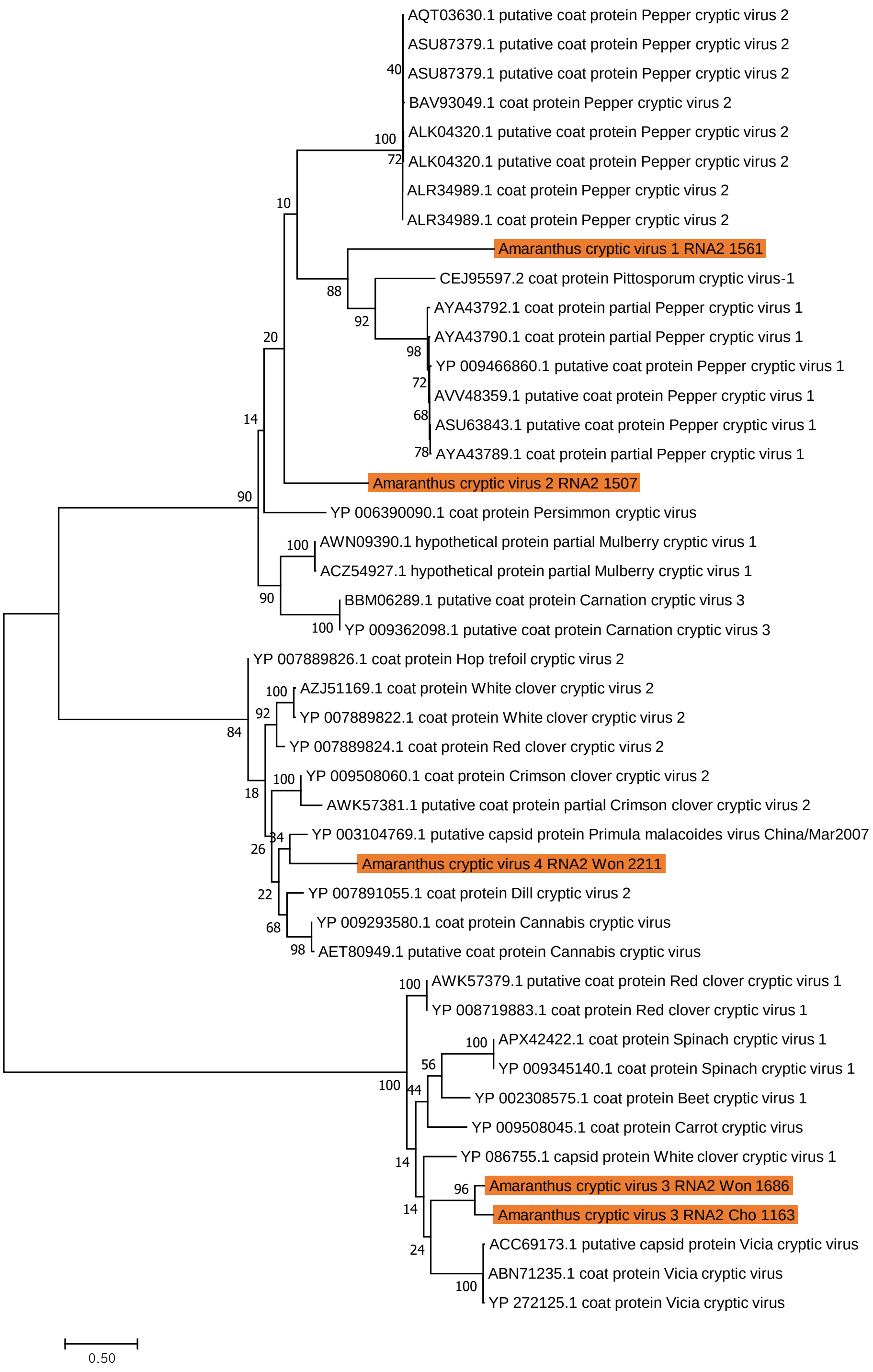

### Ambrosia cryptic virus 1–2 using RdRp

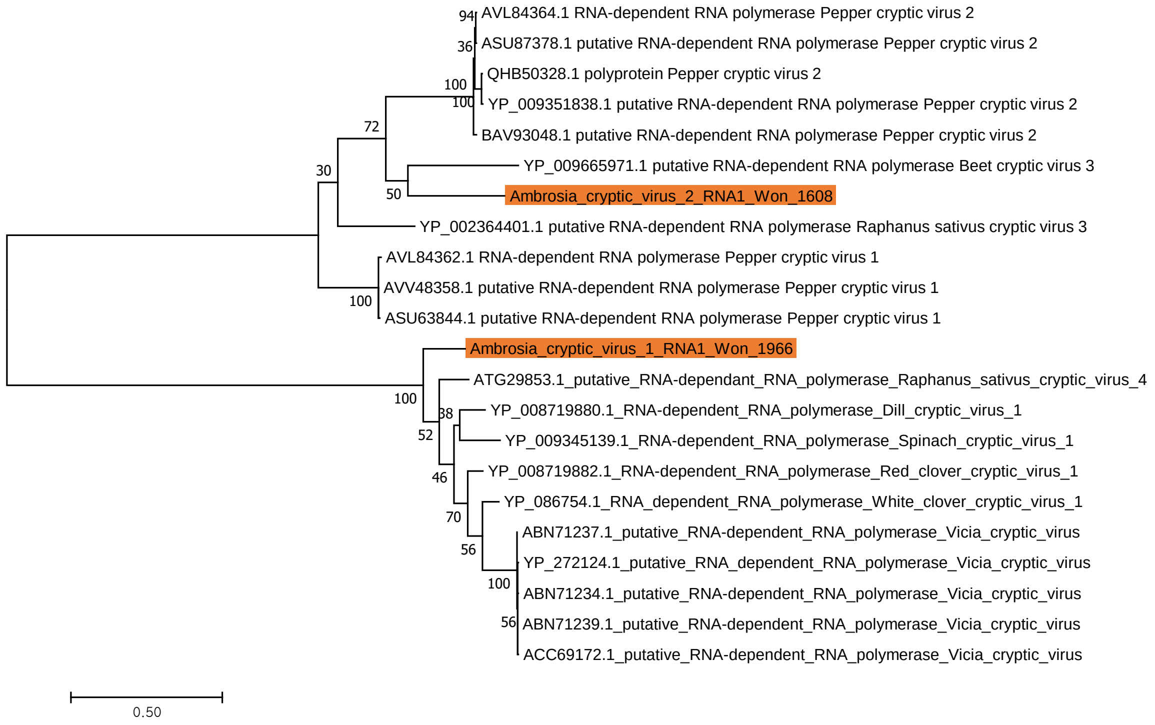

### Ambrosia cryptic virus 1–2 using CP

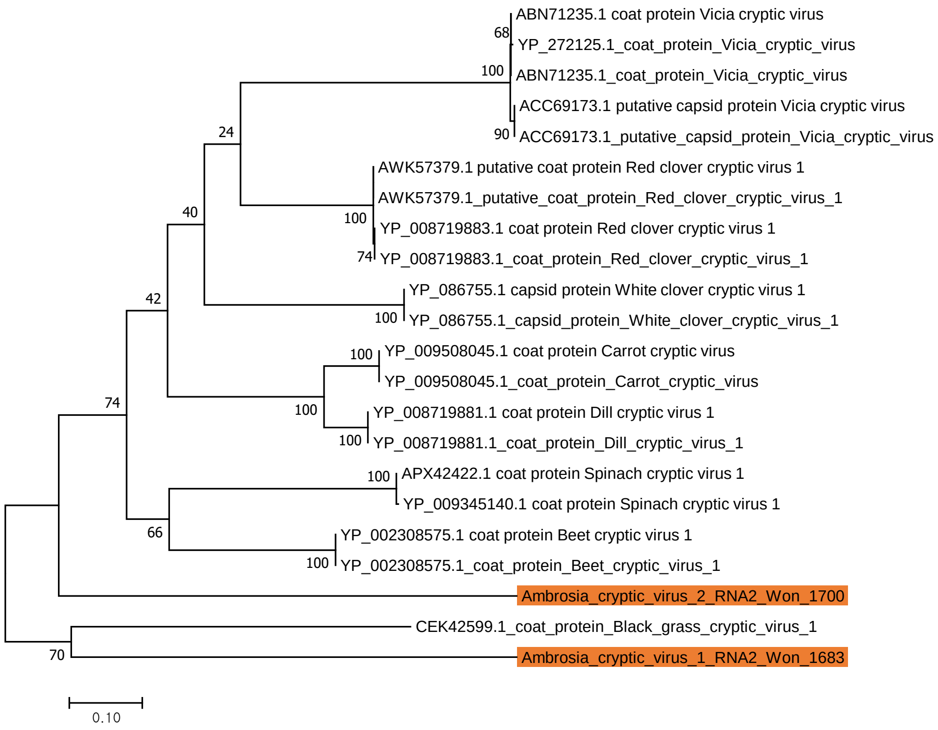

### Camellia cryptic virus 1–2 using RdRp

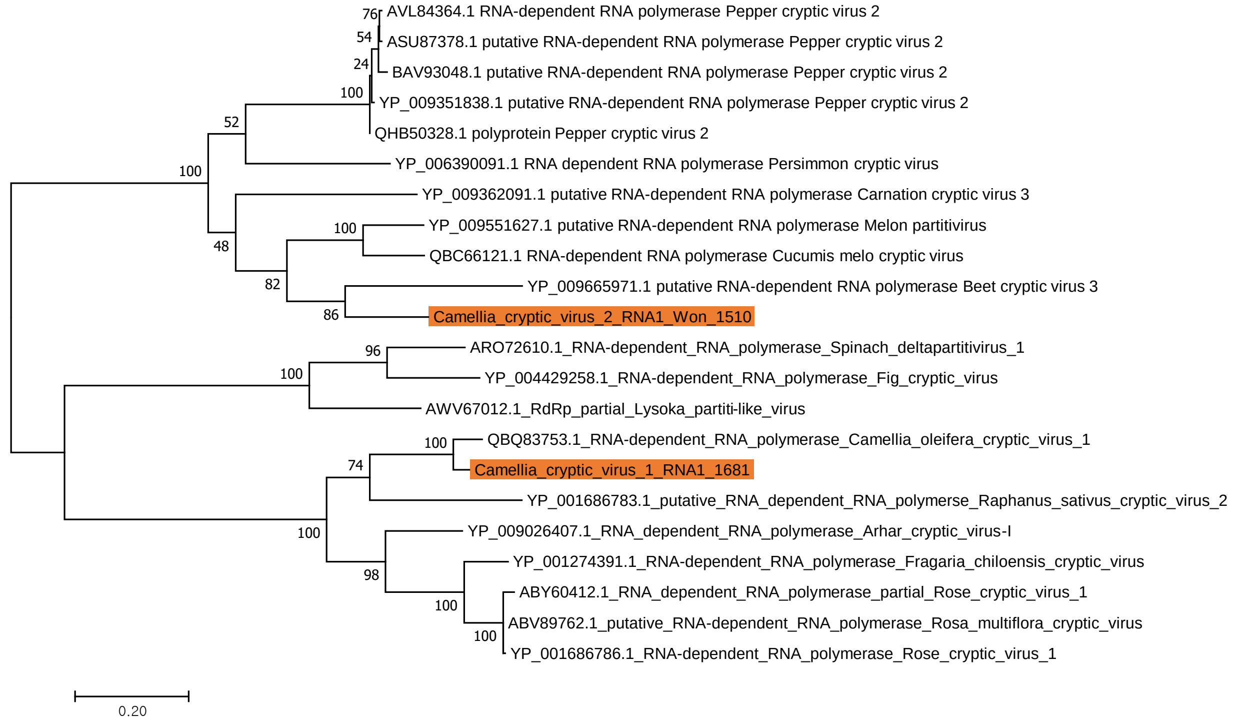

### Camellia cryptic virus 1–2 using CP

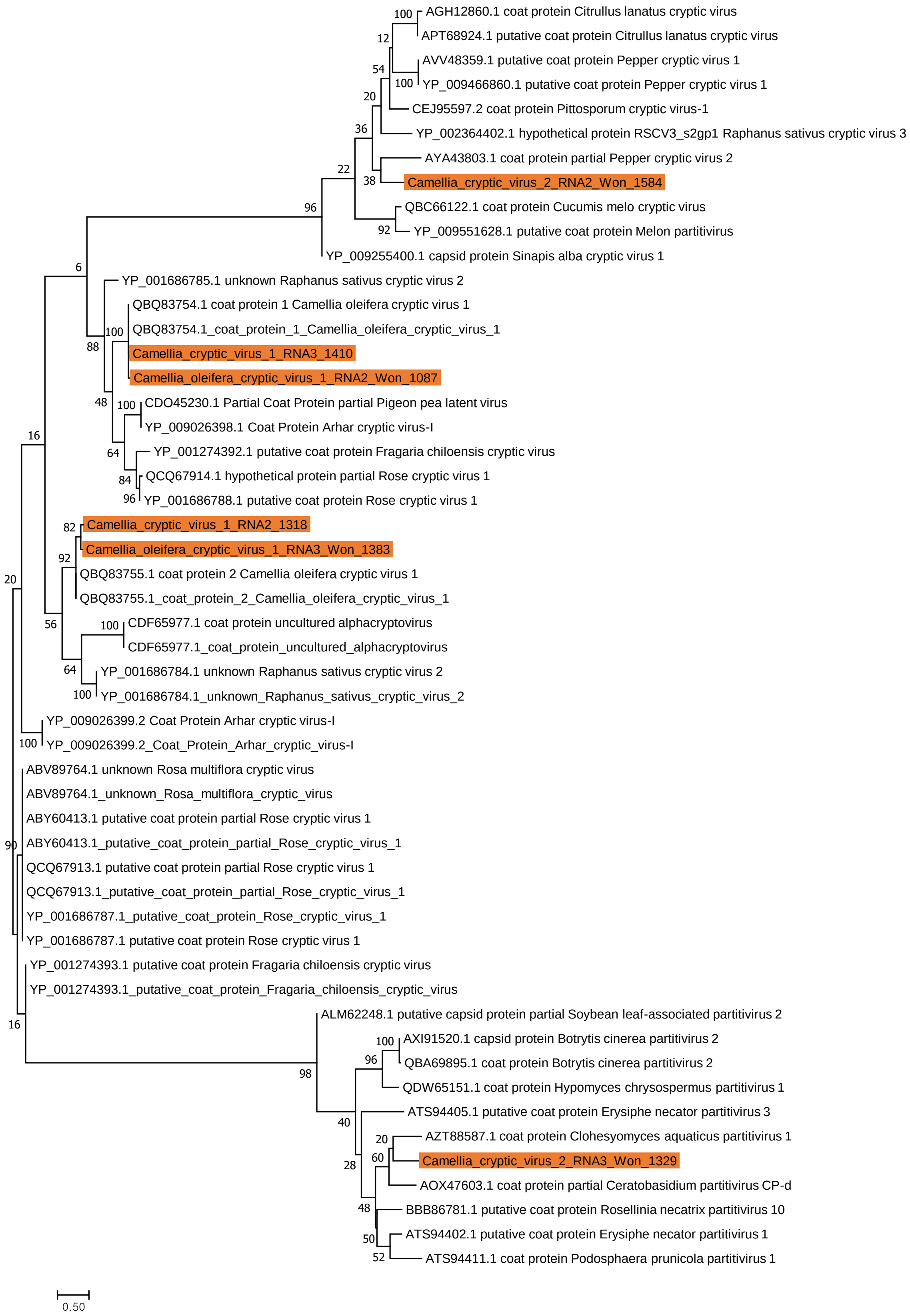

### Dactylorhiza cryptic virus 1–3 using RdRp

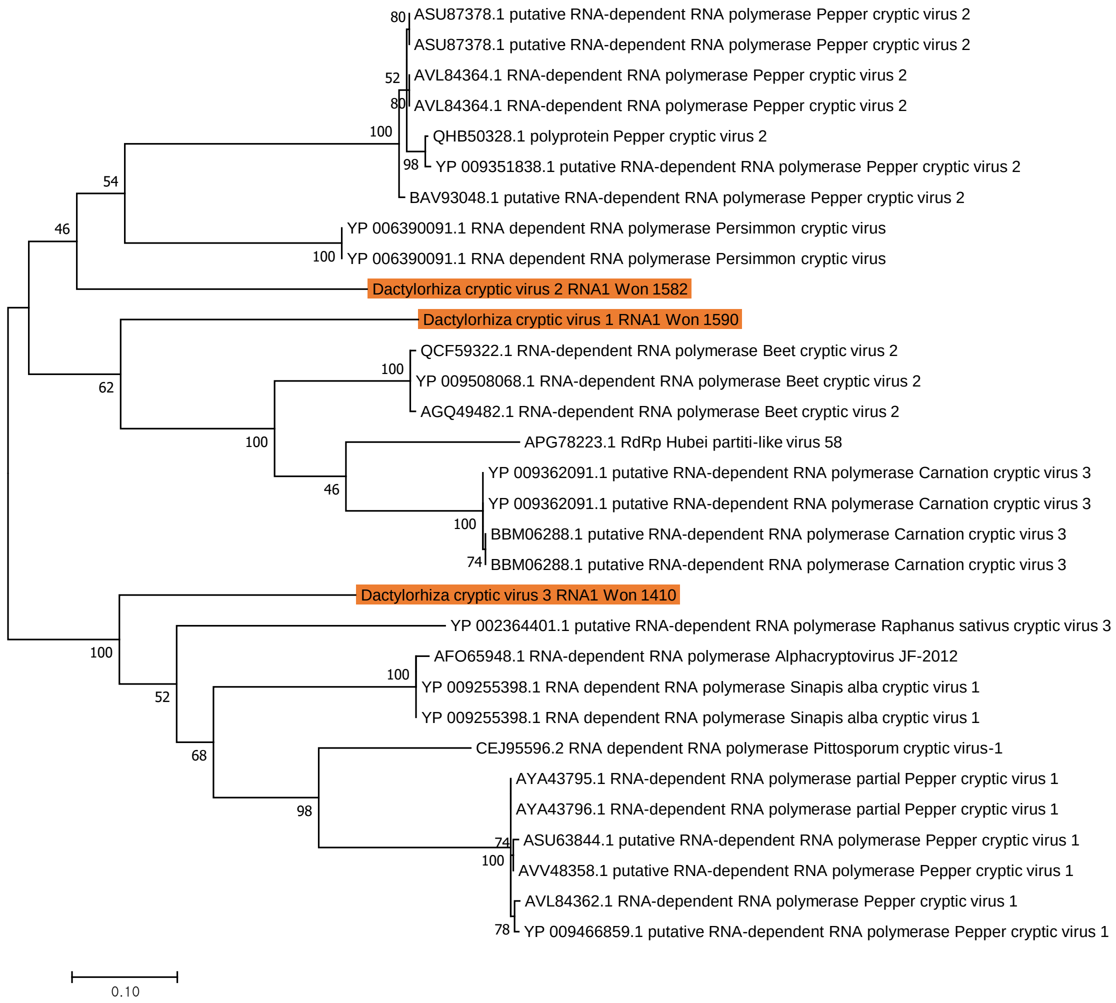

### Dactylorhiza cryptic virus 1–3 using CP

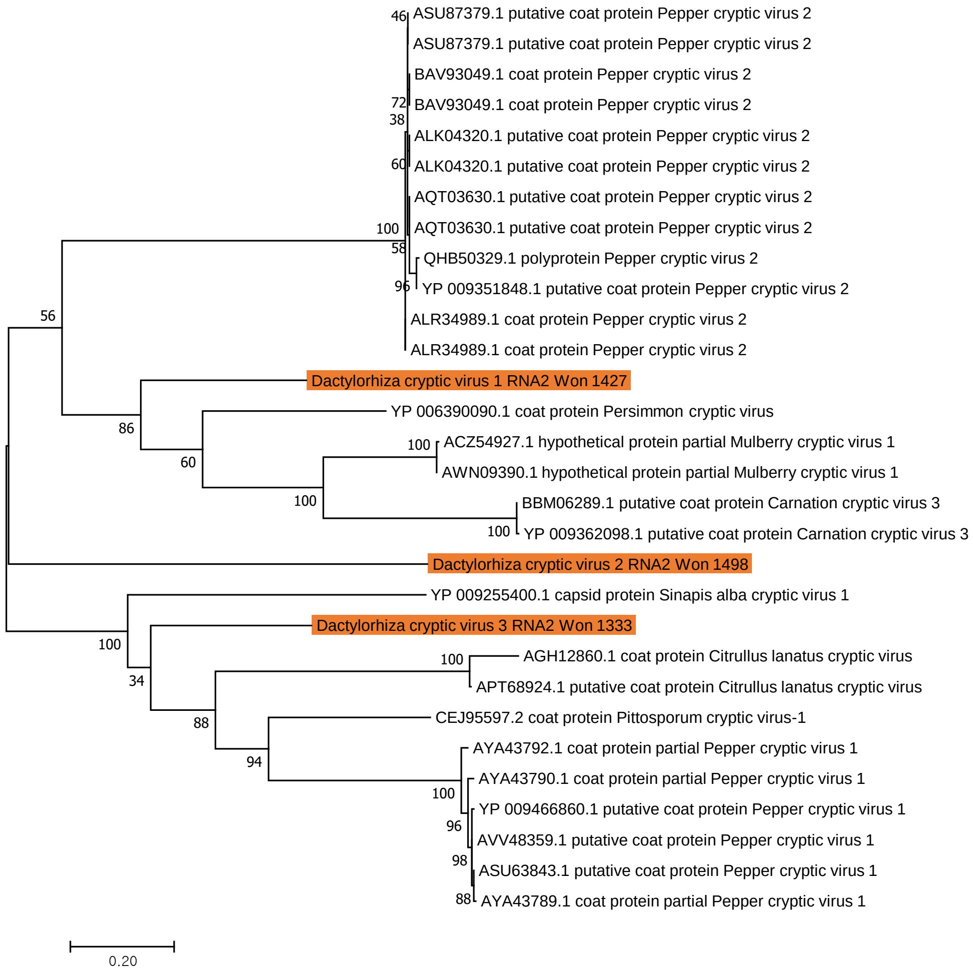

### Lomandra cryptic virus 1 using RdRp

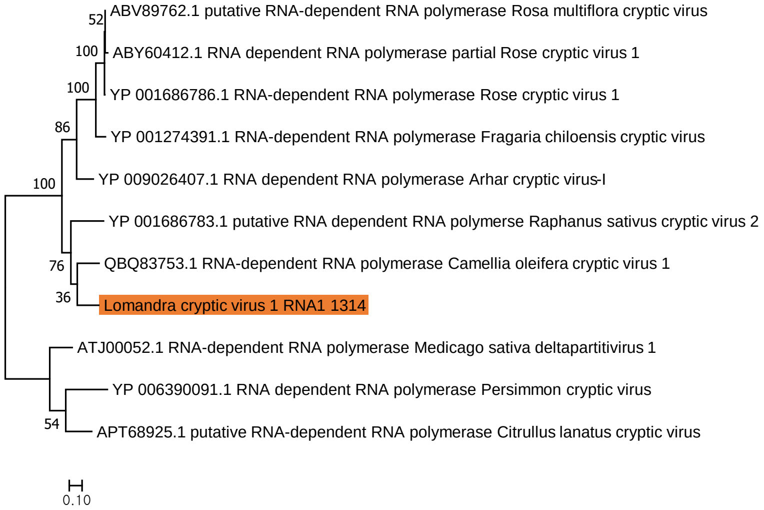

### Lomandra cryptic virus 1 using CP

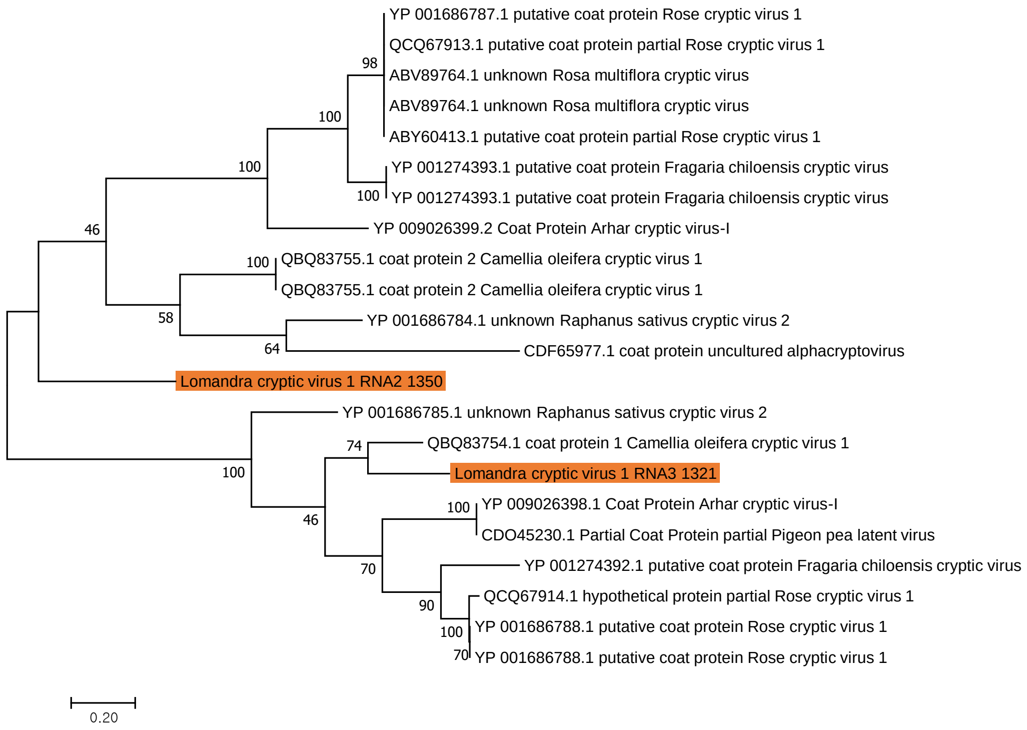

### Helianthus cryptic virus 1 using RdRp

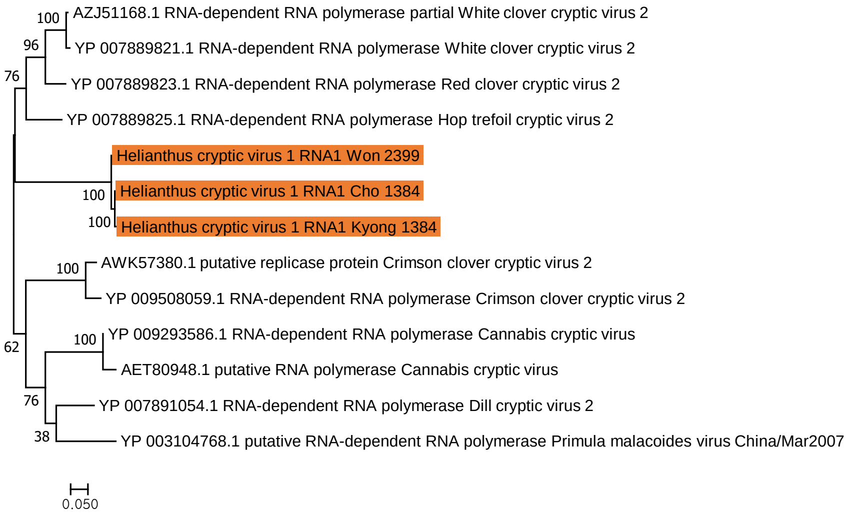

### Helianthus cryptic virus 1 using CP

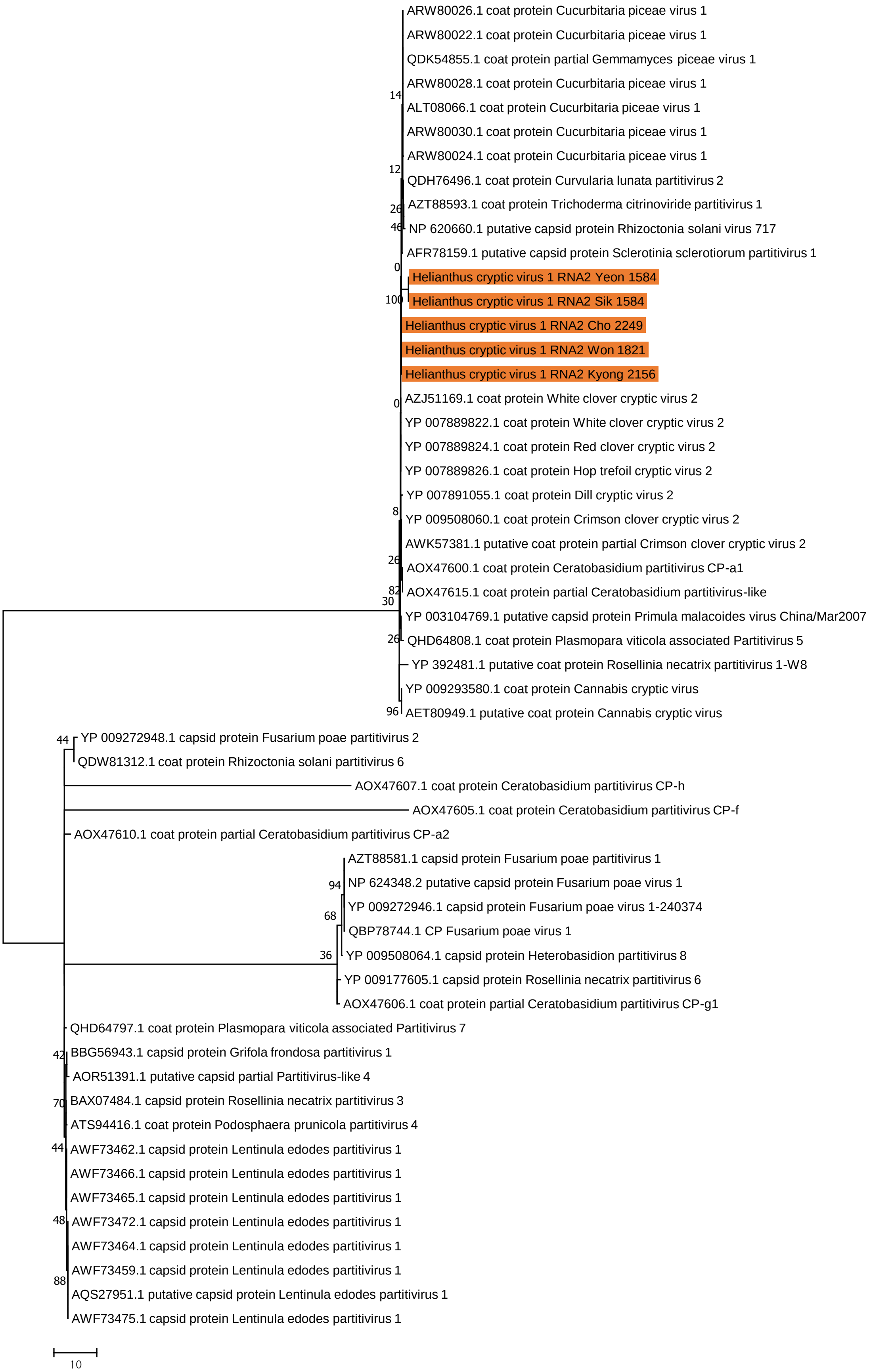

### Panax cryptic virus 1–4 using RdRp

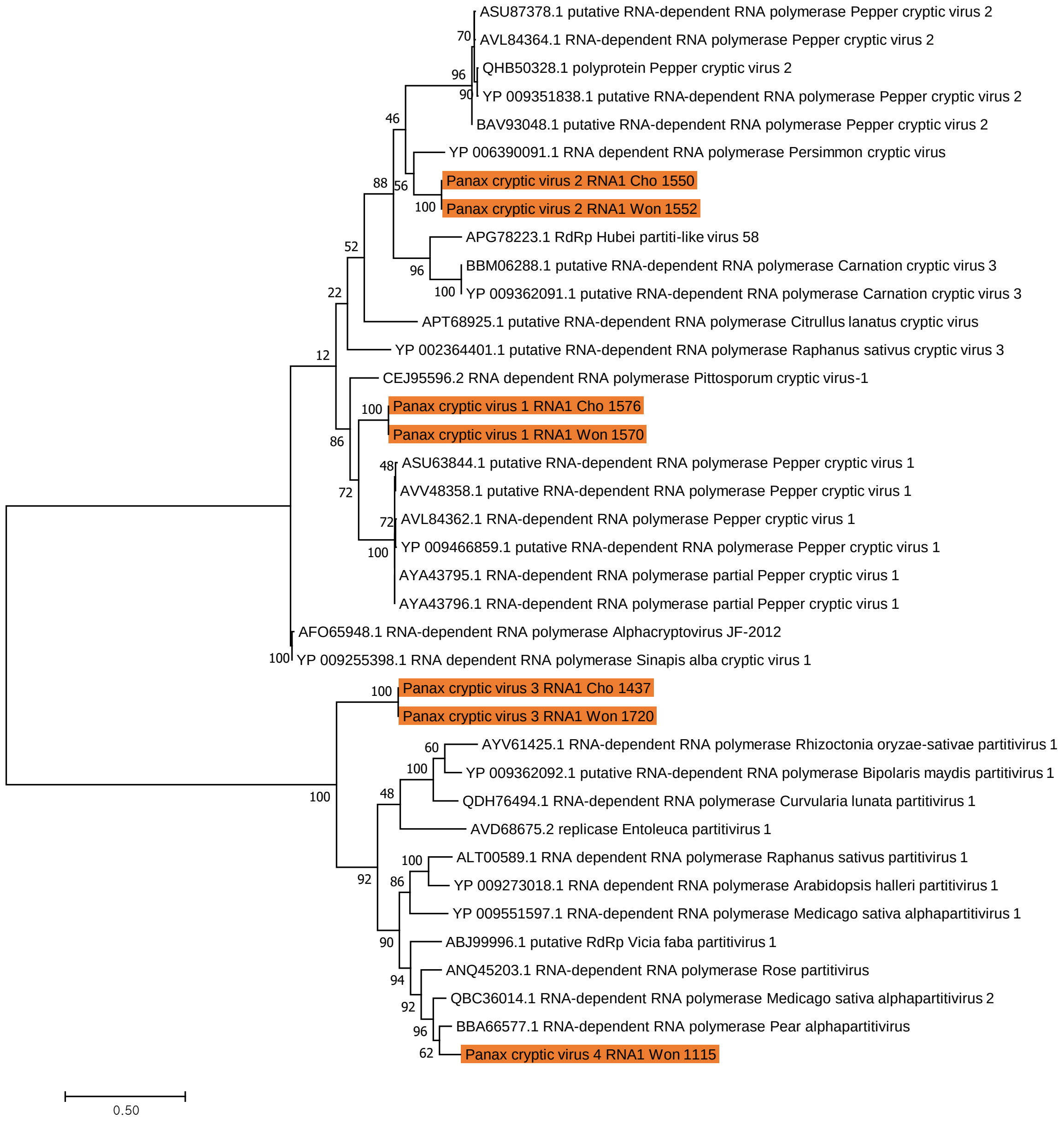

### Panax cryptic virus 1–4 using CP

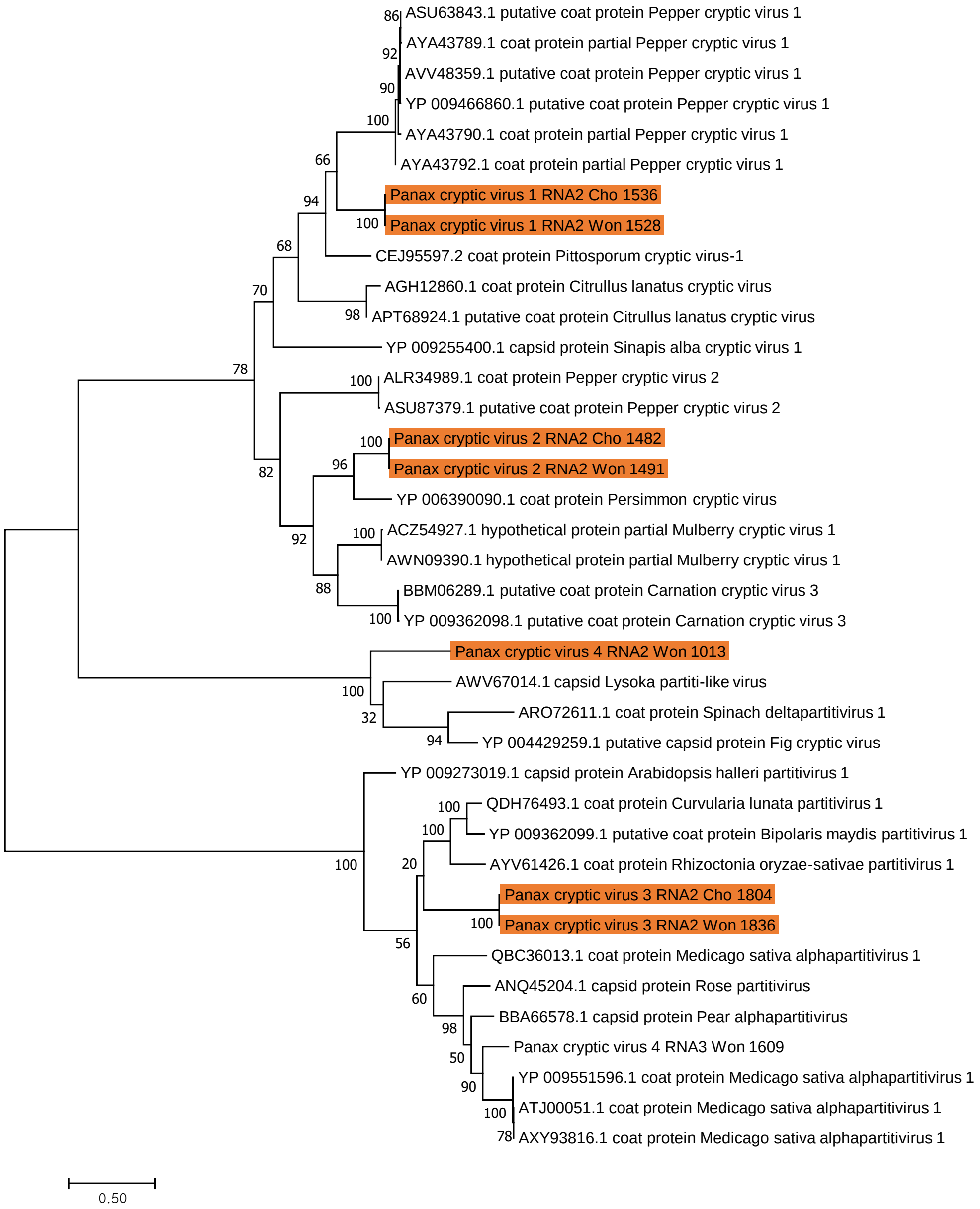

### Rhodiola cryptic virus 1–2 using RdRp

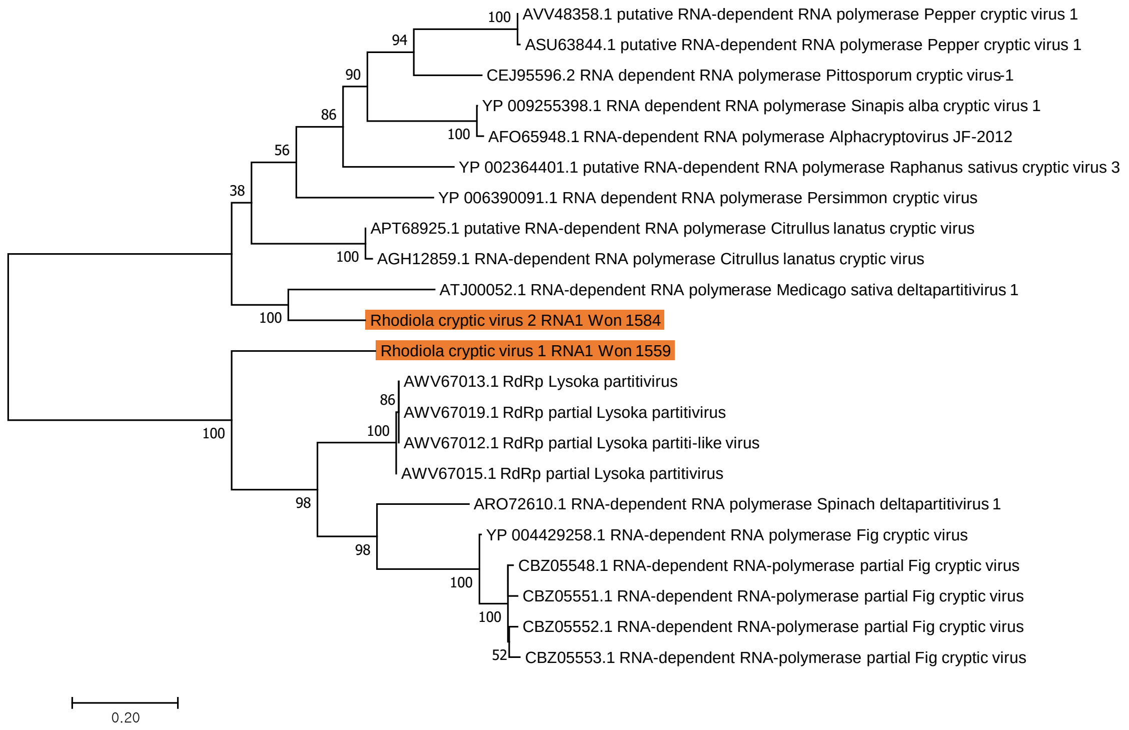

### Rhodiola cryptic virus 1–2 using CP

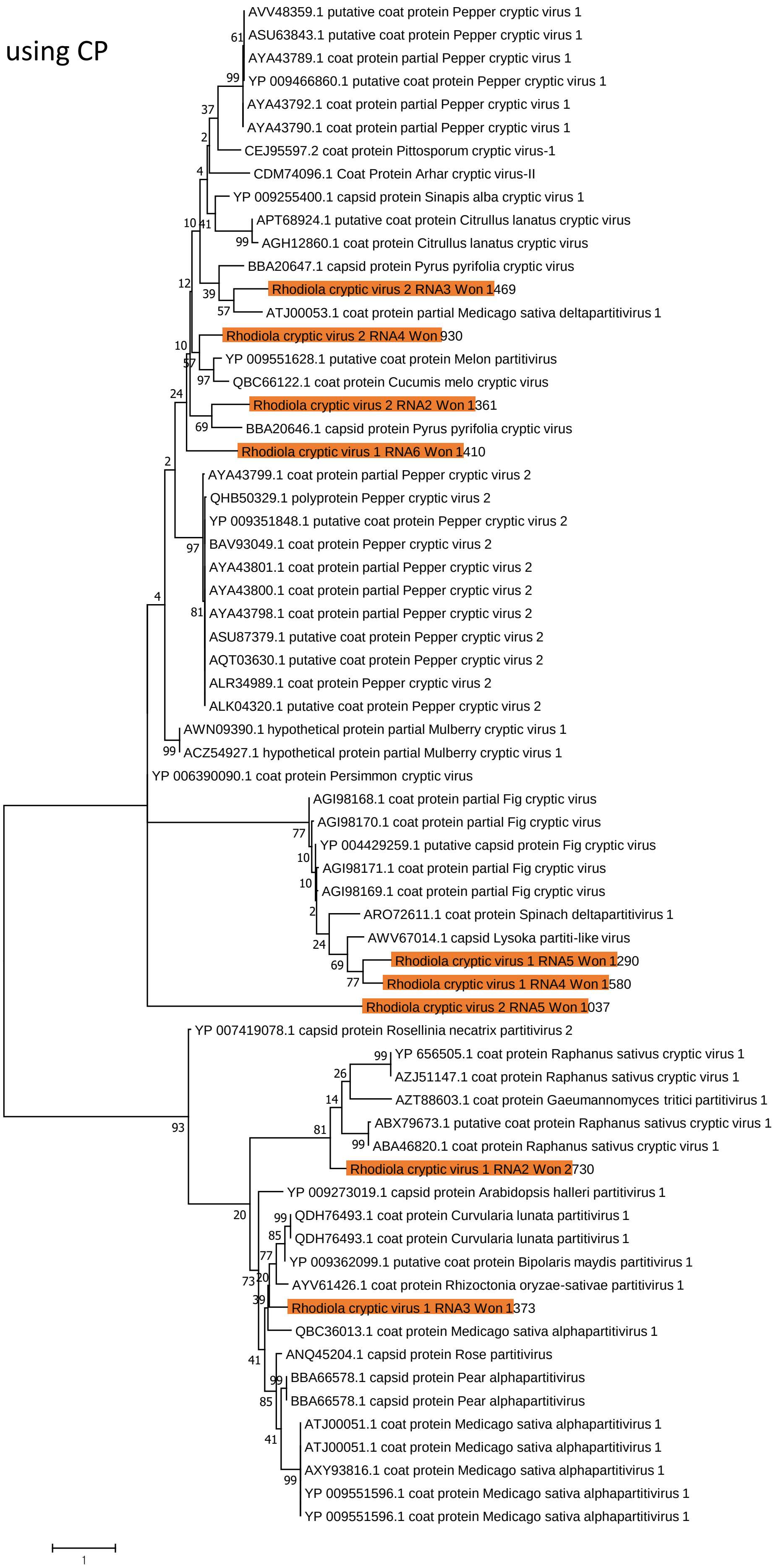

### Vicia cryptic virus using RdRp

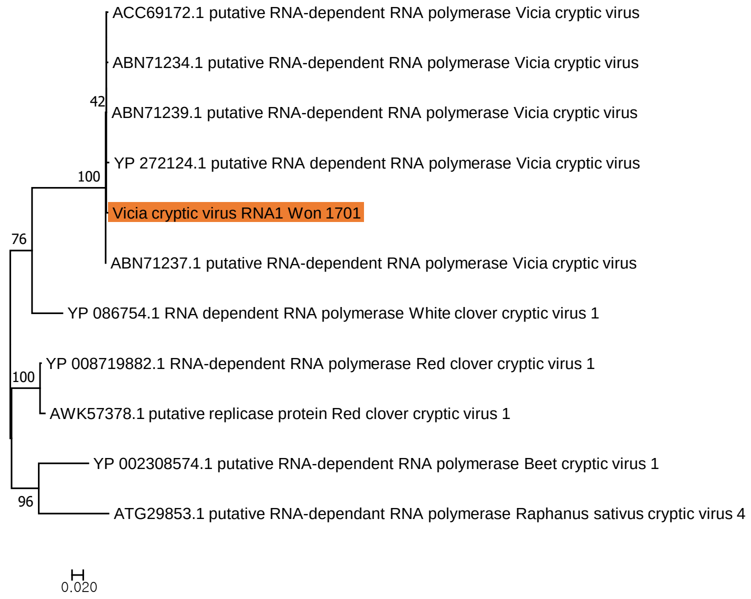

### Vicia cryptic virus  using CP

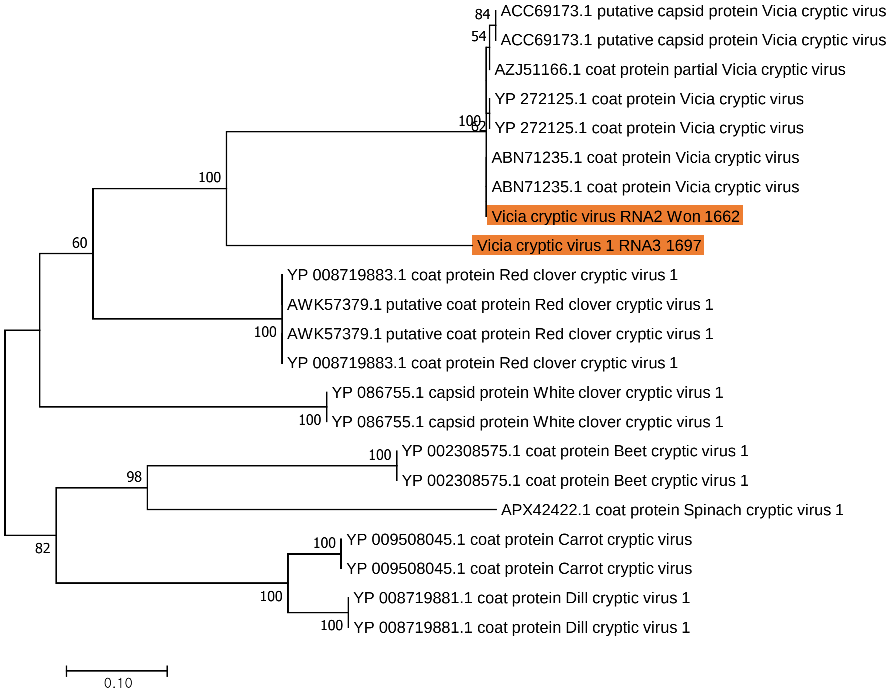
